# Tetrodotoxin-sensitive sodium channels mediate action potential firing and excitability in menthol-sensitive Vglut3-lineage sensory neurons

**DOI:** 10.1101/670620

**Authors:** Theanne N. Griffith, Trevor A. Docter, Ellen A. Lumpkin

## Abstract

Small-diameter vesicular glutamate transporter 3-lineage (Vglut3^lineage^) dorsal root ganglion (DRG) neurons play an important role in mechanosensation and thermal hypersensitivity; however, little is known about their intrinsic electrical properties. We therefore set out to investigate mechanisms of excitability within this population. Calcium microfluorimetry analysis of male and female mouse DRG neurons demonstrated that the cooling compound menthol selectively activates a subset of Vglut3^lineage^ neurons. Whole-cell recordings showed that small-diameter Vglut3^lineage^ DRG neurons fire menthol-evoked action potentials and exhibited robust, transient receptor potential melastatin 8 (TRPM8)-dependent discharges at room temperature. This heightened excitability was confirmed by current-clamp and action potential phase-plot analyses, which showed menthol-sensitive Vglut3^lineage^ neurons to have more depolarized membrane potentials, lower firing thresholds, and higher evoked firing frequencies compared with menthol-insensitive Vglut3^lineage^ neurons. A biophysical analysis revealed voltage-gated sodium channel (Na_V_) currents in menthol-sensitive Vglut3^lineage^ neurons were resistant to entry into slow inactivation compared with menthol-insensitive neurons. Multiplex *in situ* hybridization showed similar distributions of tetrodotoxin (TTX)-sensitive NaVs transcripts between TRPM8-positive and -negative Vglut3^lineage^ neurons; however, Na_V_1.8 transcripts, which encode TTX-resistant channels, were more prevalent in TRPM8-negative neurons. Conversely, pharmacological analyses identified distinct functional contributions of Na_V_ subunits, with Na_V_1.1 driving firing in menthol-sensitive neurons, whereas other small-diameter Vglut3^lineage^ neurons rely primarily on TTX-resistant NaV channels. Additionally, when Na_V_1.1 channels were blocked, the remaining Na_V_ currents readily entered into slow inactivation in menthol-sensitive Vglut3^lineage^ neurons. Thus, these data demonstrate that TTX-sensitive NaVs drive action potential firing in menthol-sensitive sensory neurons and contribute to their heightened excitability.

**Significance Statement:** Somatosensensory neurons encode various sensory modalities including thermoreception, mechanoreception, nociception and itch. This report identifies a previously unknown requirement for tetrodotoxin-sensitive sodium channels in action potential firing in a discrete subpopulation of small-diameter sensory neurons that are activated by the cooling agent menthol. Together, our results provide a mechanistic understanding of factors that control intrinsic excitability in functionally distinct subsets of peripheral neurons. Furthermore, as menthol has been used for centuries as an analgesic and anti-pruritic, these findings support the viability of Na_V_1.1 as a therapeutic target for sensory disorders.

## Introduction

Small-diameter dorsal root ganglion (DRG) neurons are sensory neurons that encode a diverse array of somatic sensations, including various forms of pain, thermosensation, itch and touch (Dubin and Patapoutian, 2010; McGlone and Reilly, 2010; Schepers and Ringkamp, 2010; Bautista et al., 2014; Liljencrantz and Olausson, 2014). This functional diversity is encompassed by small-diameter DRG neurons of the vesicular glutamate transporter 3 lineage (Vglut3^lineage^), which comprise approximately 15% of DRG neurons (Lou et al., 2013). For example, a subpopulation of Vglut3^lineage^ neurons are unmyelinated, low threshold mechanoreceptors (C-LTMRs) that encode tactile stimuli (Seal et al., 2009). Furthermore, transient receptor potential melastatin (TRPM8)-expressing Vglut3^lineage^ neurons are proposed to mediate oxaliplatin-induced cold hypersensitivity (Draxler et al., 2014). The diverse physiological processes in which these neurons have been implicated suggests they engage distinct transduction mechanisms to encode sensory information. Yet, the molecular determinants involved in transmitting electrical signals in discrete subpopulations of Vglut3^lineage^ neurons remain poorly understood.

Following membrane depolarization, activation of voltage-gated sodium channels (Na_V_s) initiate action potentials. In sensory neurons, both action potential shape and discharge frequency transmit important information (Djouhri et al., 1998; Park and Dunlap, 1998; Liu et al., 2017), a concept that is clearly illustrated in small-diameter nociceptors. These neurons predominantly express tetrodotoxin (TTX)-sensitive Na_V_1.7 channels, as well as TTX-resistant Na_V_1.8 and Na_V_1.9 subunits. Many small-diameter, nociceptive DRG neurons exhibit a prominent TTX-resistant sodium current that produces a “shoulder” during action potential repolarization, therefore increasing action potential duration (Ritter and Mendell, 1992; Djouhri et al., 1998; Blair and Bean, 2002). The inactivation kinetics of TTX-resistant sodium currents during this shoulder likely allow for a greater contribution of high-voltage activated calcium channels, which may increase calcium entry and could be particularly relevant to neurotransmitter release at presynaptic terminals (Blair and Bean, 2002). Additionally, the kinetics of slow inactivation of TTX-resistant voltage-gated sodium channels in nociceptive neurons controls firing rate adaption in response to sustained depolarization (Blair and Bean, 2003; Choi et al., 2007). Thus, the molecular identity and biophysical properties of Na_V_s expressed in a given neuron impacts action potential firing and sensory coding.

Despite the contributions of small-diameter Vglut3^lineage^ neurons to somatosensation, the Na_V_s that mediate action potential firing in these neurons remain unknown. Such information can provide important insights as to how developmentally related, yet functionally diverse, DRG neurons differentially engage Na_V_s to transmit sensory information. Accordingly, we asked whether subpopulations of small-diameter Vglut3^lineage^ DRG neurons possess measurable differences in intrinsic excitability and, if so, whether such differences reflect the contributions of functionally distinct Na_V_ subunits. Here, we show that small-diameter Vglut3^lineage^ neurons activated by the cooling compound menthol exhibit heightened excitability compared with menthol-insensitive neurons, firing robust trains of TRPM8-dependent action potentials at room temperature. Furthermore, *in vitro* electrophysiological and pharmacological analyses revealed that unlike nociceptors, TTX-sensitive ion channels including Na_V_1.1, drive action potential firing and mediate excitability in these neurons.

## Materials & Methods

### Key Resources

**Table 1.** contains a list of this study’s key resources, suppliers and unique identifying information. RRIDs are provided for mouse strains and antibodies.

| REAGENT or RESOURCE | SOURCE | IDENTIFIER |
| --- | --- | --- |
| **Antibodies** |  |  |
| Rabbit polyclonal anti-DsRed | Clontech Laboratories | Cat# 632496,<br>RRID:AB_10013483 |
| Goat anti Rabbit polyclonal Alexa-594 | Thermo Fisher Scientific | Cat# A-11037,<br>RRID:AB_2534095 |
| **Chemicals, Peptides, and Recombinant Proteins** |  |  |
| Collagenase Type P | Sigma-Aldrich | Cat# 11249002001 |
| O.C.T. Compound | Tissue-Tek | Cat# 4583 |
| Trypsin-EDTA 0.05% | Fisher Scientific | Cat# 25300054 |
| Fluoromount-G | Thermo Fisher Scientific | Cat# 00-4958-02 |
| Laminin | Sigma-Aldrich | Cat# L2020-1MG |
| MEM | Fisher Scientific | Cat# 11098050 |
| Penicillin-Streptomycin | Fisher Scientific | Cat# 15-140-122 |
| MEM vitamin solution | Thermo Fisher Scientific | Cat# SH3059901 |
| B-27 supplement | Fisher Scientific | Cat# 17504044 |
| Horse serum, heat inactivated | Thermo Fisher Scientific | Cat# 26050070 |
| Fura-2, AM | Thermo Fisher Scientific | Cat# F1221 |
| Pluronic F-127 | Thermo Fisher Scientific | Cat# P3000MP |
| Human Na<sub>v</sub>1.7 plasmid | Dr. Manu Ben-Johny |  |
| DMEM | Thermo Fisher Scientific | Cat# 11995073 |
| Fetal Bovine Serum | Thermo Fisher Scientific | Cat# A3840101 |
| Tetrodotoxin | Abcam | Cat# ab120054 |
| 4,9-anhydro-tetrodotoxin | R&D Systems | Cat# 6159/100U |
| PF 05089771 | Tocris | Cat# 5931 |
| ICA 121431 | Tocris | Cat# 5066/10 |
| PN3a | Deuis et al., 2017 |  |
| **Critical Commercial Assays** |  |  |
| Plasmid Maxi Kit | Qiagen | Cat# 12162 |
| RNAscope Fluorescent Multiplex kit | Advanced Cell Diagnostics | Cat# 320850 |
| **Deposited Data** |  |  |
| Source Data (freely available through Mendeley) | This paper | http://dx.doi.org/10.17632/bxsvmy2zh3.1 |
| Movie of calcium imaging experiment | This paper | http://dx.doi.org/10.17632/bxsvmy2zh3.1 |
| **Experimental Models: Organisms/Strains** |  |  |
| Mouse: *Slc17a8*<sup>iCre</sup>; Tg(Slc17a8-icre)1Edw/SealJ | The Jackson Laboratory | RRID:IMSR_JAX:018147 |
| Mouse: B6.Cg-Gt(ROSA)26Sor<sup>tm14(CAG-tdTomato)Hze/J</sup> | The Jackson Laboratory | RRID:IMSR_JAX:007914 |
| Homo sapiens: embryonic kidney cells stably transfected with Na<sub>v</sub>1.1, β1 and β2 subunits | (Kahlig et al., 2010) |  |
| Homo sapiens: embryonic kidney cells stably transfected with Na<sub>v</sub>1.6 | Dr. Lori Isom |  |
| **Oligonucleotide** |  |  |
| *Trpm8* probe channel 3 | Advanced Cell Diagnostics | Cat# 420451-C3 |
| *Scn1a* probe channel 2 | Advanced Cell Diagnostics | Cat# 434181-C2 |
| *Scn8a* probe channel 2 | Advanced Cell Diagnostics | Cat# 434191-C2 |
| *Scn9a* probe channel 2 | Advanced Cell Diagnostics | Cat# 313341-C2 |
| *Scn10a* probe channel 2 | Advanced Cell Diagnostics | Cat# 426011-C2 |
| **Software and Algorithms** |  |  |
| pCLAMP 10 Software Suite | Molecular Devices | https://www.moleculardevices.com |
| Image J | (Schneider et al., 2012) | https://imagej.nih.gov/ij/index.html |
| Matlab 2017b | MathWorks | www.mathworks.com |
| Software, algorithm, custom (MatLab) | This paper | https://github.com/thelumpkinlab/Calcium-Imaging-Analysis-Script |
| Metamorph v7.5.6 | Molecular Devices | https://www.moleculardevices.com |
| Metafluor v7.6.3 | Molecular Devices | https://www.moleculardevices.com |
| Prism v7.0b | GraphPad | www.graphpad.com |

### Animals

Animal use was conducted according to guidelines from the National Institutes of Health’s Guide for the Care and Use of Laboratory Animals and was approved by the Institutional Animal Care and Use Committee of Columbia University Medical Center. Mice were maintained on a 12 h light/dark cycle, and food and water was provided ad libitum. *Slc17a8^iCre^*(stock #018147, Grimes et al., 2011) and *Rosa26^Ai14^* mice (stock #007914, Madisen et al., 2010) were obtained from Jackson Laboratories and bred to produce *Slc17a8^iCre^;Rosa26^Ai14^*mice. Genotyping was performed through Transnetyx. Adult *Slc17a8^iCre^;Rosa26^Ai14^*mice (4-16 weeks old) of either sex were used for all experiments.

### DRG culture preparation

DRGs were harvested from *Slc17a8^iCre^;Rosa26^Ai14^* mice and transferred to Ca^2+^-free, Mg^2+^-free Hank’s solution (HBSS, Gibco) containing the following (in mM): 137.9 NaCl, 5.3 KCl, 0.34 Na_2_HPO_4_, 0.44 K_2_HPO_4_, 5.6 glucose, 4.2 NaHCO_3_, 0.01% phenol red. Processes were trimmed to reduce the amount of plated non-neuronal cells. Ganglia were treated with collagenase (1.5 mg/ml, Type P, Sigma-Aldrich) in HBSS for 20 min at 37°C followed by 0.05% Trypsin-EDTA (Gibco) for 3 min with gentle rotation. Trypsin was neutralized with culture media (MEM, with L-glutamine, Phenol Red, without sodium pyruvate) supplemented with 10% horse serum (heat-inactivated, Gibco), 10 U/ml penicillin, 10 μg/ml streptomycin, MEM vitamin solution (Gibco), and B-27 supplement. Serum-containing media was decanted and cells were triturated using a fire-polished Pasteur pipette in a serum-free MEM culture media containing the supplements listed above. Cells were plated on laminin-treated (0.05 mg/ml) glass coverslips, which had previously been washed in 2N NaOH for at least 4 h, rinsed with 70% ethanol and UV-sterilized. Cells were then incubated at 37°C in 5% CO_2_. Cells were used for electrophysiological experiments 16-24 h after plating.

### HEK cell culture and transfection

Stably transfected HEK293 cell lines expressing human Na_V_1.1 (Kahlig et al., 2010), Na_V_1.6 (Dr. Lori Isom, University of Michigan), or HEK293 cells transiently transfected with a cDNA construct containing human Na_V_1.7 (Dr. Manu Ben-Johny, Columbia University) were used. The Na_V_1.7 plasmid was sequenced following transformation and extraction (Genewiz, see Mendeley dataset). HEK cells were grown in DMEM (GIBCO 11995) containing 10% FBS (Thermo Fisher Scientific A3840101), 1% Penicillin-Streptomycin (Thermo Fisher Scientific 15-140-122). Media for HEK cells stably expressing Na_V_1.1 or Na_V_1.6 also contained 400 μg/ml G418 (Fisher Scientific 10-131-035) to select for transfected cells. A calcium phosphate protocol was used to co-transfect Na_V_1.7 and green fluorescent protein into HEK 293 cells. Briefly, 2 M CaCl_2_, cDNAs, and sterile water were mixed together and added dropwise to a 2x solution of hepes buffered saline. The final solution was added dropwise to HEK293 cells that were plated the day before on glass cover slips coated with 0.05 mg/ml laminin. Cells were incubated at 37°C in 5% CO_2_ with the transfection solution for 3 h, followed by two washes with sterile phosphate buffered saline and addition of new cell culture media. Electrophysiological recordings were performed 24-72 hours post transfection.

### Electrophysiology

Whole-cell voltage- and current-clamp recordings made from small-diameter (capacitance ≤ 25 pF), TdTomato-expressing (Vglut3^lineage^) dissociated DRG neurons and HEK cells were performed with patch pipettes pulled from standard borosilicate glass (1B150F-4, World Precision Instruments) with a P-97 puller (Sutter Instruments). For neuronal recordings, patch pipettes had resistances of 3-6 MΩ when filled with an internal solution containing the following (in mM): 120 K-methylsulfonate, 10 KCl, 10 NaCl, 5 EGTA, 0.5 CaCl_2_, 10 HEPES, 2.5 MgATP, pH 7.2 with KOH, osmolarity 300 mOsm. For HEK cell recordings, patch pipettes had resistances of 1.5-3 MΩ when filled with an internal solution containing the following (in mM): 140 CsF, 10 NaCl, 2 MgCl_2_, 0.1 CaCl_2_, 1.1 EGTA, 10 HEPES, pH 7.2 with CsOH, osmolarity ∼310 mOsm. Seals and whole-cell configuration were obtained in an external solution containing (in mM): 145 NaCl, 5 KCl, 10 HEPES, 10 glucose, 2 CaCl_2_, 2 MgCl^2^, pH 7.3 with NaOH, osmolarity ∼320 mOsm. Series resistance was compensated by 80%. In experiments with DRG neurons where currents from voltage-gated sodium channels were recorded, after the whole-cell configuration was established and neurons were tested for sensitivity to menthol, a modified external solution was applied (in mM): 105 NaCl, 40 TEA-Cl, 10 HEPES, 2 BaCl_2_, 13 glucose, 0.03 CdCl_2_, pH 7.3 with NaOH, osmolarity ∼320 mOsm. This modified solution was used in HEK cell recordings following seal acquisition in a standard external solution. All solutions used were allowed to warm to ambient temperature before each experiment to ensure all recordings were made at room temperature (20°C-23°C). After each experiment, the recording chamber was thoroughly cleaned with Milli-Q water. For experiments using Na_V_ inhibitors, drugs were applied to neurons for 1 min in current-clamp mode in the absence of injected current; therefore, cells were recorded at their intrinsic V_m_.

### Data acquisition and analysis

Currents and voltages were acquired and analyzed using pClamp software (version 10, Molecular Devices). Recordings were obtained using an Axopatch 200b patch-clamp amplifier and a Digidata 1440A, and filtered at 5 kHz and digitized at 10 kHz. Analysis was performed using Clampfit 10 (Molecular Devices). All voltages were corrected for the measured liquid junction potential (−7 mV) between internal and external recording solutions. Phase plots were constructed from the first derivative of the somatic membrane potential (dV/dT) versus the instantaneous somatic membrane potential. Action potential threshold was calculated as the membrane potential at which the phase plot slope reached 10 mV ms^-1^ (Kress et al., 2008; Yu et al., 2008). Duration at the base was calculated by measuring the duration of the action potential starting at the resting membrane potential (V_m_) and ending when the repolarization phase again passes the initial V_m_. Following determination of menthol sensitivity in gap-free recording mode, some cells were not further analyzed using phase plot analysis due to low digital gain during recordings (n=3) or deteriorating cell health (n=2).

### Pharmacology

TTX was from Abcam. PF 05089771 and ICA 121431 were from Tocris. PN3a was a generous gift from Dr. Irina Vetter (Institute for Molecular Biosciences, University of Queensland). All other chemicals were from Sigma-Aldrich.

### Calcium imaging

Dissociated DRG neurons were loaded for 45 min with 10 mM Fura-2AM (Invitrogen), supplemented with 0.01% Pluronic F-127 (wt/vol, Invitrogen), in external solution. Images were acquired using MetaMorph software (version 7) and displayed as the ratio of 340 nm to 380 nm. Neurons were identified by eliciting calcium responses with a high potassium solution (140 mM) at the end of each experiment. Neurons were considered sensitive to an agonist if the average ratio during the 30 s following agonist application was ≥ 15% above baseline. Image analysis was performed using custom MATLAB scripts.

### Multiplex in situ hybridization

DRG sections cut at 25-µm thickness were processed for RNA *in situ* detection using an RNAscope Fluorescent Detection Kit according to manufacturer’s instructions (Advanced Cell Diagnostics, Hayward, California, USA) with the following modifications: upon harvesting, DRG were fixed in 4% paraformaldehyde for 15 min and then incubated in 30% sucrose for 2 h at 4°C. DRG were embedded in Optimal Cutting Temperature Compound (Sakura) and stored at - 80°C until sectioned. The following RNAscope probes were used: *Trpm8* (420451-C3, mouse), *Scn1a* (434181-C2, mouse), *Scn8a* (434191-C2, mouse), *Scn9a* (313341-C1, mouse), and *Scn10a* (426011-C2, mouse). *In situ* hybridization was followed by incubation at 4°C overnight with a rabbit anti-dsRed (1:3000, Clontech: 632475) primary antibody. Sections were then incubated at room temperature for 1 h with a goat anti-rabbit AlexaFluor 594-conjugated secondary antibody (Thermo Fisher Scientific, A-11012). Samples were mounted with Fluoromount-G (Fisher Scientific). Specimens were imaged in three dimensions (1-µm axial steps) on a Zeiss Exciter confocal microscope (LSM 5) equipped with a 40X, 1.3 NA objective lens. Images were analyzed using ImageJ software. Neurons considered positive for a given Na_V_ subunit had signal equal to or greater than one standard deviation above background.

### Experimental design and statistical analysis

Summary data are presented as mean ± standard deviation from *n* cells. For quantitative analysis of *in situ* hybridization data, at least three biological replicates per condition were used and the investigator was blinded to Na_V_ subunit for analysis. Statistical differences between menthol-sensitive and menthol-insensitive populations were assessed using an unpaired Student’s *t* test (two-tailed) for normally distributed datasets. A Mann Whitney test was used for populations that did not conform to Gaussian distributions or had different variances. To estimate IC_50_ values for Na_V_ antagonists, inhibitor versus response curves were fit with the following relation: Normalized current=100/(1+[Inhibitor]/IC_50_). Kinetic data were fit with single or double-exponential relations. The voltage-dependence of slow inactivation was fit with the Boltzmann equation: Fraction available=Minimum+[(Maximum-Minimum)/(1+exp(*V_50_*-Vm)/*k*)], where *V_50_* denotes the membrane potential at which half the channels are inactivated and *k* denotes the Boltzmann constant/slope factor. Differences between fits were assessed with an Extra sum-of-squares F test. Statistical tests and fit parameters are listed in the *Results* and/or figure legends. Statistical significance in each case is denoted as follows: \**P* < 0.05, \*\**P* < 0.01; \*\*\**P* < 0.001, and \*\*\*\**P* < 0.0001. Statistical tests and curve fits were performed using Prism 7.0 (GraphPad Software).

## Results

### Menthol-sensitivity is restricted to Vglut3^lineage^ DRG neurons

Vglut3^lineage^ sensory neurons are a heterogeneous population. To identify functionally distinct subpopulations within this group, we tested the responsiveness of Vglut3^lineage^ neurons to capsaicin, chloroquine and menthol (Figure 1, Figure 1-1), which activate nociceptors, pruritoceptors and cold receptors, respectively. We performed calcium microfluorimetry while applying various chemosensory stimuli to acutely cultured DRG neurons (<24 h) harvested from adult male and female *Slc17a8^iCre^;Rosa26^Ai14^* mice. In these mice, neurons that express Vglut3 at any point during development are labeled with a TdTomato fluorescent reporter (Figure 1A). Neurons were identified by robust calcium responses to high-potassium depolarization (784 total DRG neurons, 331 Vglut3^lineage^, 453 non-Vglut3^lineage^, n=3 mice; Figure 1 B–D). Approximately 8% of Vglut3^lineage^ neurons were activated by the TRPM8 agonist menthol, whereas no non-Vglut3^lineage^ neurons responded to the compound (Figure 1E). Conversely, both populations contained neurons that were activated by the TRP vanilloid 1 (TRPV1) agonist capsaicin; however, comparatively fewer Vglut3^lineage^ neurons were capsaicin-sensitive compared with non-Vglut3^lineage^ neurons (∼9% vs. ∼46%, respectively). Few neurons of either group responded to both menthol and capsaicin (5/322 Vglut3^lineage^ neurons), or to chloroquine (1/322 Vglut3^lineage^ neurons and 4/460 non-Vglut3^lineage^ neurons), a pruritogen that signals through MrgprA3 and TRP ankyrin 1 (TRPA1; Wilson et al., 2011). Interestingly, a comparison of un-normalized baseline fura-2 ratios showed that menthol-sensitive neurons had slightly elevated baseline calcium signals compared to menthol-insensitive neurons (F_340_/F_380_ = 0.52 ± 0.05 vs. 0.45 ± 0.03, n = 5 coverslips, *P* = 0.029, unpaired Student’s *t* test, two-tailed, Figure 1F). This analysis builds upon prior work (Draxler et al., 2014) by demonstrating that menthol sensitivity is restricted to the Vglut3^lineage^ population.

**Figure 1.**
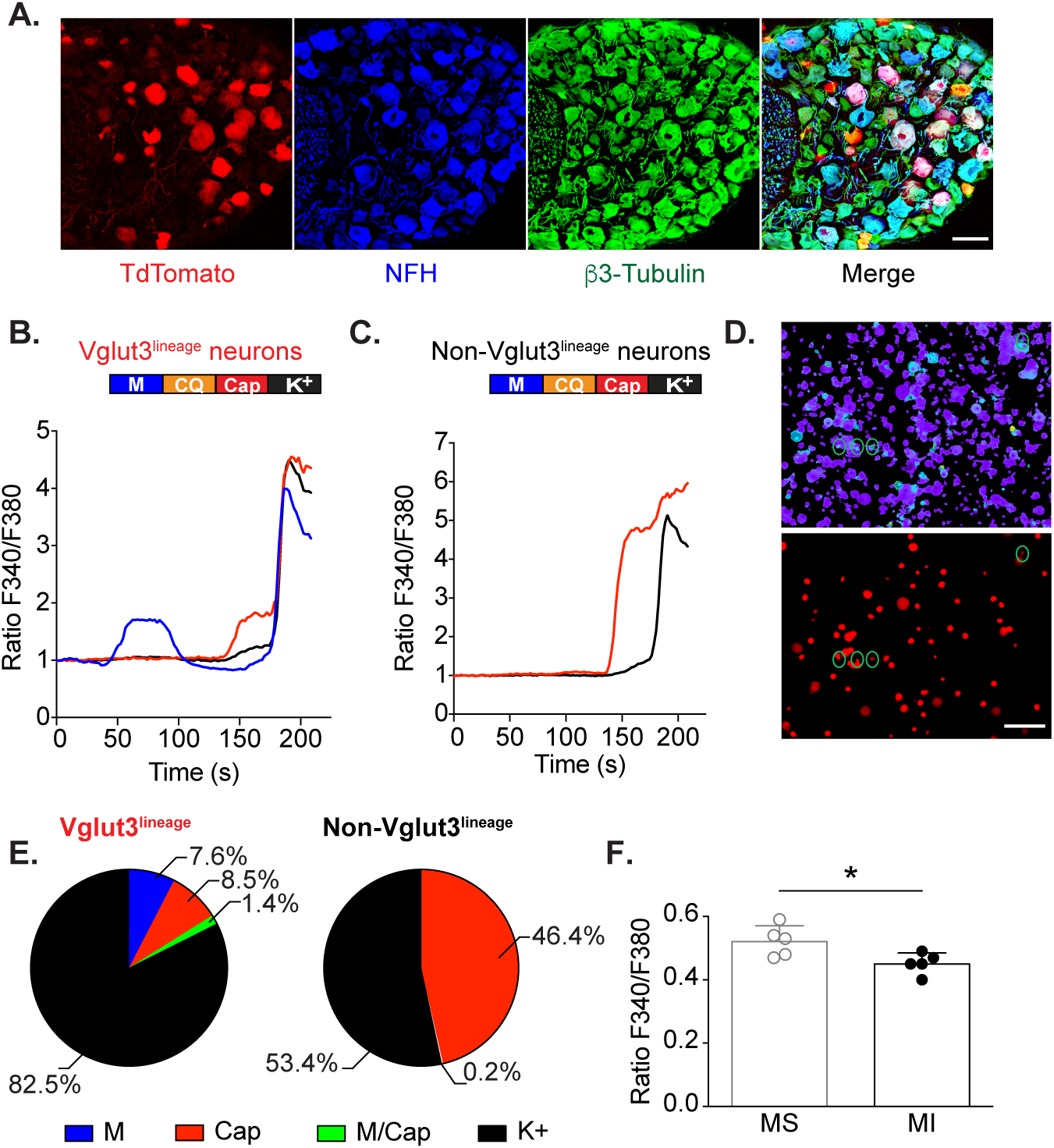
Menthol-sensitivity is restricted to Vglut3^lineage^ DRG neurons. **A.** Representative confocal images of DRG sections (25 μm) from adult *Slc17a8^iCre^;Rosa26^Ai1^* mice immunostained with anti-DsRed (TdTomato, red), anti-Neurofilament Heavy (NFH, blue), and anti-β3-Tubulin (green). Images were acquired using a 20x, 0.8 NA air objective (**B-C).** Baseline normalized, representative traces depicting Fura2-AM ratio (F340/F380) versus time traces of averaged responses from Vglut3^lineage^ (**B**) and non-Vglut3^lineage^ (**C**) DRG neurons to various chemosensory stimuli (menthol 100 μM [M, green trace], chloroquine,1 mM [CQ], capsaicin 1 μM [Cap, red trace], and high K+ Ringer’s [K+, black trace]). Colored bar indicates time of agonist application. **D**. Images of calcium transients in live, dissociated *Slc17a8^iCre^;Rosa26^Ai1^* DRG neurons quantified in **B** and **C**. (Top) Fura2-AM calcium microfluorimetry following menthol application. Green circles indicate menthol-sensitive DRG neurons. (Bottom) Fluorescent image showing TdTomato-expressing (Vglut3^lineage^) DRG neurons. **E.** Quantification of percentage of Vglut3^lineage^ (n = 331) and non-Vglut3^lineage^ (n = 453) neurons responding to individual agonists. **F.** Quantification of baseline calcium signals between Vglut3^lineage^ menthol-sensitive neurons and menthol-insensitive neurons (both Vglut3^lineage^ and non-Vglut3^lineage^). Significance was determined using an unpaired Student’s *t* test; *\*P < 0.05*. Data represented as mean ± S.D. Scale bars, 100 μm.

The majority of menthol-sensitive DRG neurons have small somata and give rise to unmyelinated axons (Takashima et al., 2007; Dhaka et al., 2008). Thus, we targeted small-diameter, Vglut3^lineage^ neurons with a membrane capacitance (C_m_) of ≤ 25 pF for functional analysis. Neurons that did not meet these criteria were not analyzed further by electrophysiology. Using gap-free current-clamp recordings, we asked whether these neurons fire action potentials in response to menthol application (100 μM). Half of small-diameter Vglut3^lineage^ neurons (31/62 neurons) fired trains of action potentials in response to menthol application (Figure 2A). A subset of neurons were subsequently exposed to 1 mM menthol, which activates TRPM8 ion channels but inhibits TRPA1 (Karashima et al., 2007; Xiao et al., 2008). All neurons examined showed a dose-dependent increase in menthol-evoked firing rates (Figure 2B), suggesting that menthol elicits firing through TRPM8 rather than TRPA1 in Vglut3^lineage^ DRG neurons. We noted that menthol-sensitive neurons were among the smallest DRG neurons *in vitro*, whereas menthol-insensitive neurons were more varied in size (Figure 2C). Consistent with this observation, the distribution of C_m_ among these menthol-sensitive neurons was well fit by a single Gaussian distribution (R^2^ = 0.986; 8.1 ± 2.9 pF; Figure 2D). Conversely, menthol-insensitive neurons were better fit by a double Gaussian distribution, (R^2^ = 0.818), with the two populations having means of 8.7 ± 3.4 and 20.6 ± 2.0 pF (Figure 2D). These data suggest that menthol-sensitive Vglut3^lineage^ neurons are a more homogenous subpopulation compared with menthol-insensitive Vglut3^lineage^ neurons.

**Figure 2.**
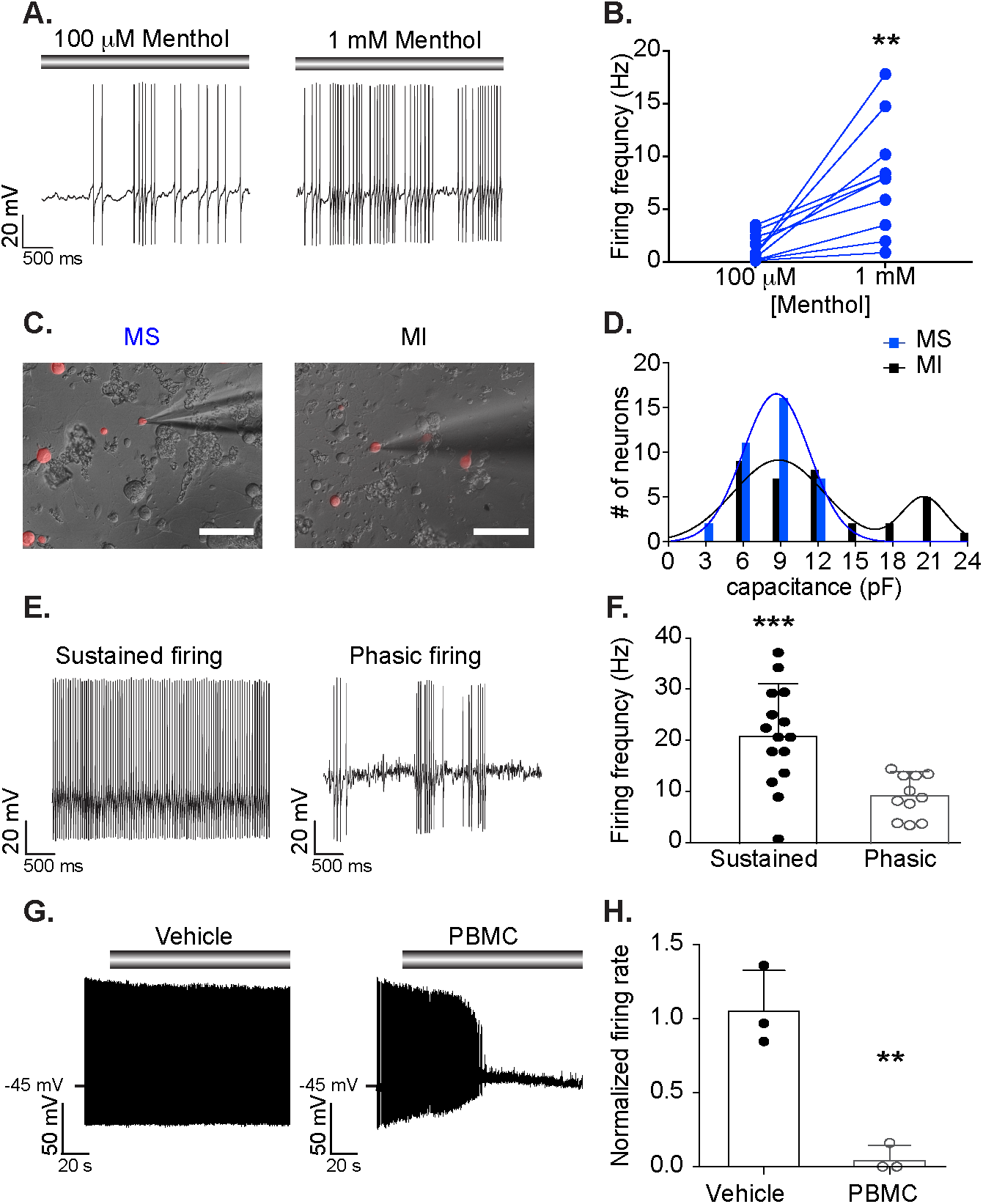
Menthol-sensitive Vglut3^lineage^ neurons fire action potential discharges at room temperature. **A.** Representative current-clamp recording from a menthol-sensitive Vglut3^lineage^ DRG neuron. Gray bar indicates menthol application (100 μM, left; 1 mM, right). **B.** Quantification of firing rates in response to 100 μM and 1 mM menthol. Lines connecting symbols indicate paired observations. Significance was determined using a paired Student’s *t* test (two-tailed); *\*\*P = 0.0036* **C.** Representative differential interference contrast image (20x, 0.75 NA air objective) of a MS (left) and a MI (right) DRG neuron in culture during electrophysiological recordings. TdTomato fluorescence indicates Vglut3^cre^ expression at some point during development (Vglut3^lineage^). **D.** Histogram of membrane capacitance measurements from MS (blue) and MI (black) Vglut3^lineage^ neurons. Lines indicate the mean(s) of the Gaussian core. **E.** Left, Representative current-clamp recording from a MS neuron exhibiting sustained firing at room temperature. Right, A different MS neuron firing with a phasic action potential discharge pattern. **F.** Quantification of average non-evoked firing frequency of MS Vglut3^lineage^ neurons at room temperature. Each individual point represents the average firing frequency of a single neuron over a five-second period. Firing frequencies for phasic-firing neurons were quantified during bursts of action potentials only. Significance was determined using an unpaired Student’s *t* test (two-tailed); *\*\*\*P = 0.0009*; **G.** Left, Representative current-clamp trace of room temperature action potential firing in a MS neuron. Vehicle treatment did not impact firing rate. Grey bar indicates vehicle application. Right, Representative current-clamp trace of inhibition of action potential firing in a MS neuron following application of the TRPM8 blocker, PBMC (25 nM). Grey bar indicates PBMC application. **H.** Quantification of relative firing rate following 90 s of vehicle or PBMC treatment. Significance was determined using an unpaired Student’s *t* test; *\*\*P 0.0*. Data represented as mean ± S.D. Scale bars, 100 μm.

### Menthol-sensitive Vglut3^lineage^ neurons fire robustly at room temperature

We next asked whether intrinsic excitability properties differed between menthol-sensitive and insensitive Vglut3^lineage^ neurons. During gap-free recordings, we noted that 87% (27/31) of menthol-sensitive Vglut3^lineage^ neurons exhibited unusually robust action potential firing prior to menthol application. Two firing patterns were observed, sustained and phasic firing (Figure 2E-F). Of the menthol-sensitive neurons that exhibited non-evoked activity, 44% exhibited phasic firing, whereas 56% maintained sustained firing during gap-free recordings. Menthol-sensitive neurons with sustained action potential discharges showed higher average firing frequencies compared with burst firing frequencies of phasic neurons (Figure 2F). By contrast, few menthol-insensitive neurons exhibited non-evoked firing during gap-free recordings (4/31), and these produced only occasional action potentials. This ongoing activity in menthol-sensitive neurons is consistent with the elevated baseline flura-2 fluorescence observed during calcium microfluorimetry experiments (Figure 1F), as well as *ex vivo* data showing sustained firing upon cold or menthol-stimulation of TRPM8-expressing DRG neuron receptive fields (Jankowski et al., 2017). Together, these results suggest that menthol-sensitive Vglut3^lineage^ population have heightened excitability under our *in vitro* recording conditions and that, within the menthol-sensitive Vglut3^lineage^ population, firing properties vary.

The ability of menthol-sensitive neurons to fire robustly at room temperature, an activating stimulus for TRPM8 (McKemy et al., 2002; Andersson et al., 2004; Tajino et al., 2011; Fujita et al., 2013; Morenilla-Palao et al., 2014; Jankowski et al., 2017; Pertusa et al., 2018), led us to ask whether this firing was dependent upon TRPM8 ion channels. We applied the selective inhibitor, PBMC (25 nM, Knowlton et al., 2011) to menthol-sensitive neurons and analyzed its effect on action potential firing at room temperature. Whereas vehicle application did not have a significant effect on firing rates, PBMC drastically reduced action potential firing at room temperature within 2 min of drug application (n = 3 neurons per group, *P* = 0.0035, unpaired Student’s *t* test, Figure 2G-H). Thus, activation of TRPM8 ion channels mediates robust ongoing action potential firing in menthol-sensitive Vglut3^lineage^ neurons.

Collectively, these data demonstrate that menthol-sensitive neurons are highly excitable subpopulation of small-diameter Vglut3^lineage^ DRG neurons, capable of maintaining sustained action potential firing *in vitro* at room temperature.

### Intrinsic excitability differs between Vglut3-lineage menthol-sensitive and menthol-insensitive neurons

To investigate the heightened intrinsic excitability in menthol-sensitive DRG neurons, we compared responses of menthol-sensitive and -insensitive Vglut3^lineage^ neurons to 500 ms current injections using phase plot analysis (Figure 3A-B). The threshold for action potential firing in menthol-sensitive neurons was significantly hyperpolarized (−28.5 ± 6.6 mV, n = 28) compared with menthol-insensitive neurons (−22.2 ± 10.4 mV, n = 31, *P* = 0.0269, unpaired Student’s *t* test; Figure 3C). Menthol-sensitive neurons also fired more action potentials in response to a current injection of 50 pA (Figure 3D). Action potential duration at the base (see Methods) was significantly shorter in menthol-sensitive neurons compared with menthol-insensitive neurons (Figure 3E). Interestingly, in gap-free recordings prior to menthol application, menthol-sensitive neurons had significantly more depolarized membrane potentials (V_m_) than menthol-insensitive neurons (−45.5 ± 4.4 mV vs. −51.2 ± 5.8 mV, n = 31 for each group, *P* < 0.0001, Mann Whitney test, Figure 3F). Thus, menthol-sensitive Vglut3^lineage^ neurons maintain a V_m_ that more closely borders action potential threshold as compared with menthol-insensitive Vglut3^lineage^ neurons. Conversely, action potential amplitude, and membrane voltage sag did not differ between the two populations (Figure 3G-H). Together, these data provide evidence that menthol-sensitive Vglut3^lineage^ neurons have more excitable membrane properties than menthol-insensitive Vglut3^lineage^ neurons.

**Figure 3.**
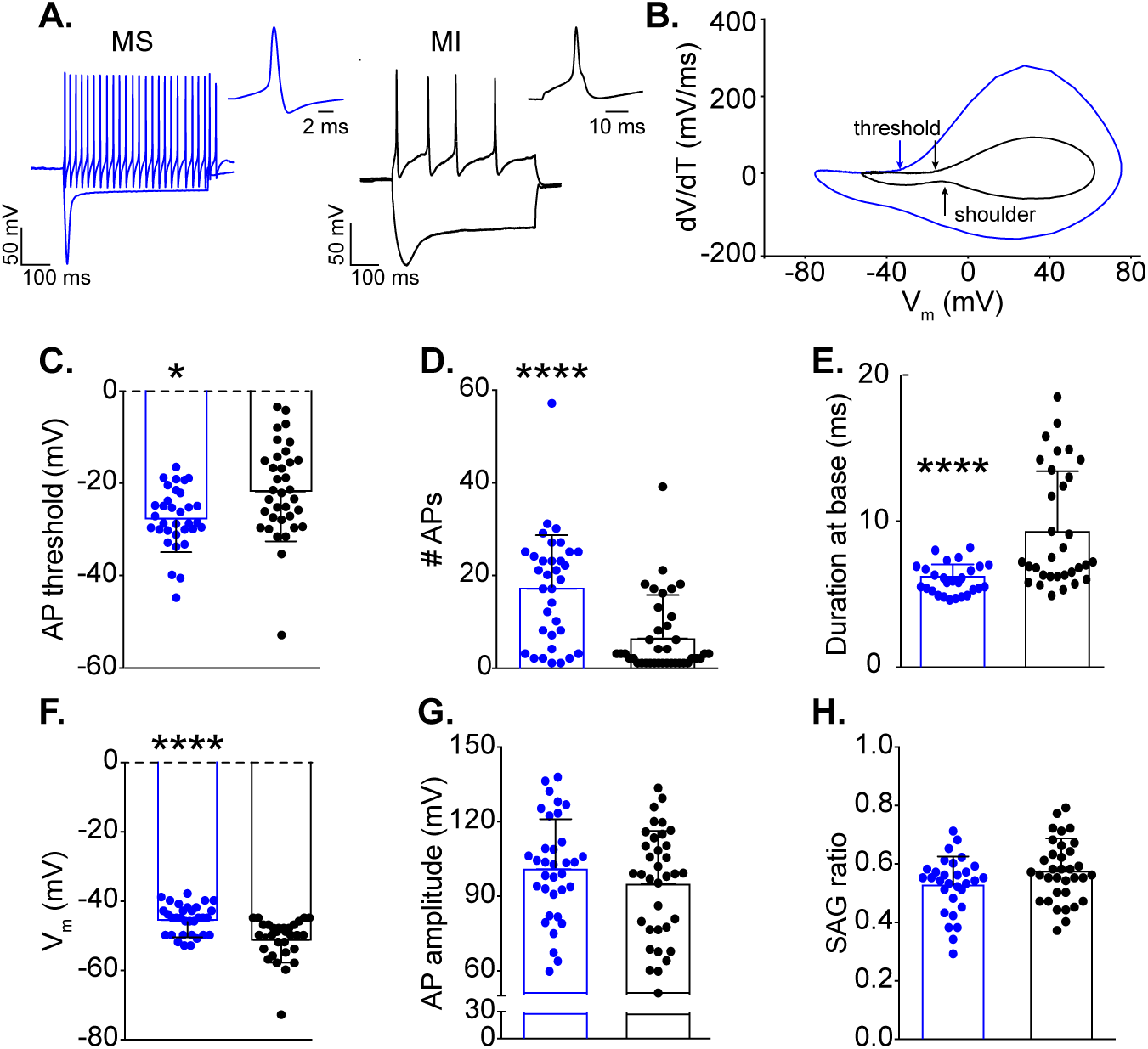
Intrinsic excitability of menthol-sensitive Vglut3^lineage^ DRG neurons. **A.** Representative current-clamp traces from a menthol-sensitive (MS, left, blue) and -insensitive (MI, right, black) Vglut3^lineage^ DRG neuron in response to −200 pA and 50 pA current injections. The single action potential for each represents the first action potential elicited by the 50 pA current injection. **B**. Phase-plots of single action potentials shown in **A**. Plots show the first derivative of the somatic membrane potential (dV/dT) versus the instantaneous somatic membrane potential. The blue curve represents the MS neuron and the black curve represents the MI neuron. Arrows indicating “threshold” are the points at which the membrane potential of the phase plot slope reached 10 mV ms^-1^. The arrow indicating “shoulder” represents the momentary slowing of membrane repolarization seen in a subpopulation of menthol-insensitive neurons. (**C-H**). Quantification of action potential threshold (**C**), number of action potentials generated in response to a 50 pA current injection (**D**), duration at base (**E**), membrane potential (**F**), half-width (**G**), and sag ratio (**H**) for MS (blue) and MI (black) Vglut3^lineage^ DRG neurons. Significance was determined using unpaired Student’s *t* tests for normally distributed populations (**C, E, G, H**) or Mann Whitney tests for non-normal distributions (**D, F**); \*\**P*<0.01, \*\*\*\**P*<0.0001. Bars denote as mean ± S.D and filled circles show data from each neuron.

Interestingly, consistent with their longer duration action potentials, 44% of menthol-insensitive neurons had a pronounced “shoulder” during the repolarization phase of the action potential (Figure 3A-B). This shoulder was completely absent in menthol-sensitive neurons. The presence of a shoulder is attributed to sodium currents mediated by TTX-resistant Na_V_1.8 and Na_V_1.9 channels (Blair and Bean, 2002). Thus, a differential contribution of Na_V_ subunits to action potential firing in these two populations of Vglut3^lineage^ neurons could underlie the observed differences in excitability.

### Differences in Na_V_ current slow inactivation kinetics in small-diameter Vglut3^lineage^ neurons

Na_V_ slow inactivation has been linked to adaptation of action potential firing in small-diameter DRG neurons, whereby sequestration of tetrodotoxin-resistant Na_V_s subunits Na_V_1.8 and Na_V_1.9 in the slow inactivated state restricts the duration of action potential discharges in response to sustained stimulation (Blair and Bean, 2003). As we found that menthol-sensitive Vglut3^lineage^ neurons are capable of maintaining prolonged action potential discharges for several minutes *in vitro* (Figure 2) and fire robustly in response to current injection (Figure 3), we hypothesized that Na_V_ slow inactivated states are unstable in menthol-sensitive Vglut3^lineage^ neurons. To test this model, we first measured Na_V_ entry into slow inactivation by delivering a conditioning pulse from −100 mV to 0 mV for 50–1600 ms between 3-ms test steps to −20 mV (Figure 4A). Consistent with our hypothesis, entry of Na_V_ currents into the slow inactive state was almost fourfold slower in menthol-sensitive neurons compared with menthol-insensitive neurons (τ = 1485 ms, n = 6 vs. τ = 376.5 ms, n = 5; *P* < 0.0001, Extra sum-of-squares F test.) Notably, after a 1600 ms conditioning pulse, ∼65% of the initial Na_V_ current in menthol-sensitive neurons was still present, whereas only ∼21% of the current remained in menthol-insensitive neurons. Thus, Na_V_ currents in menthol-sensitive neurons are resistant to slow inactivation.

**Figure 4.**
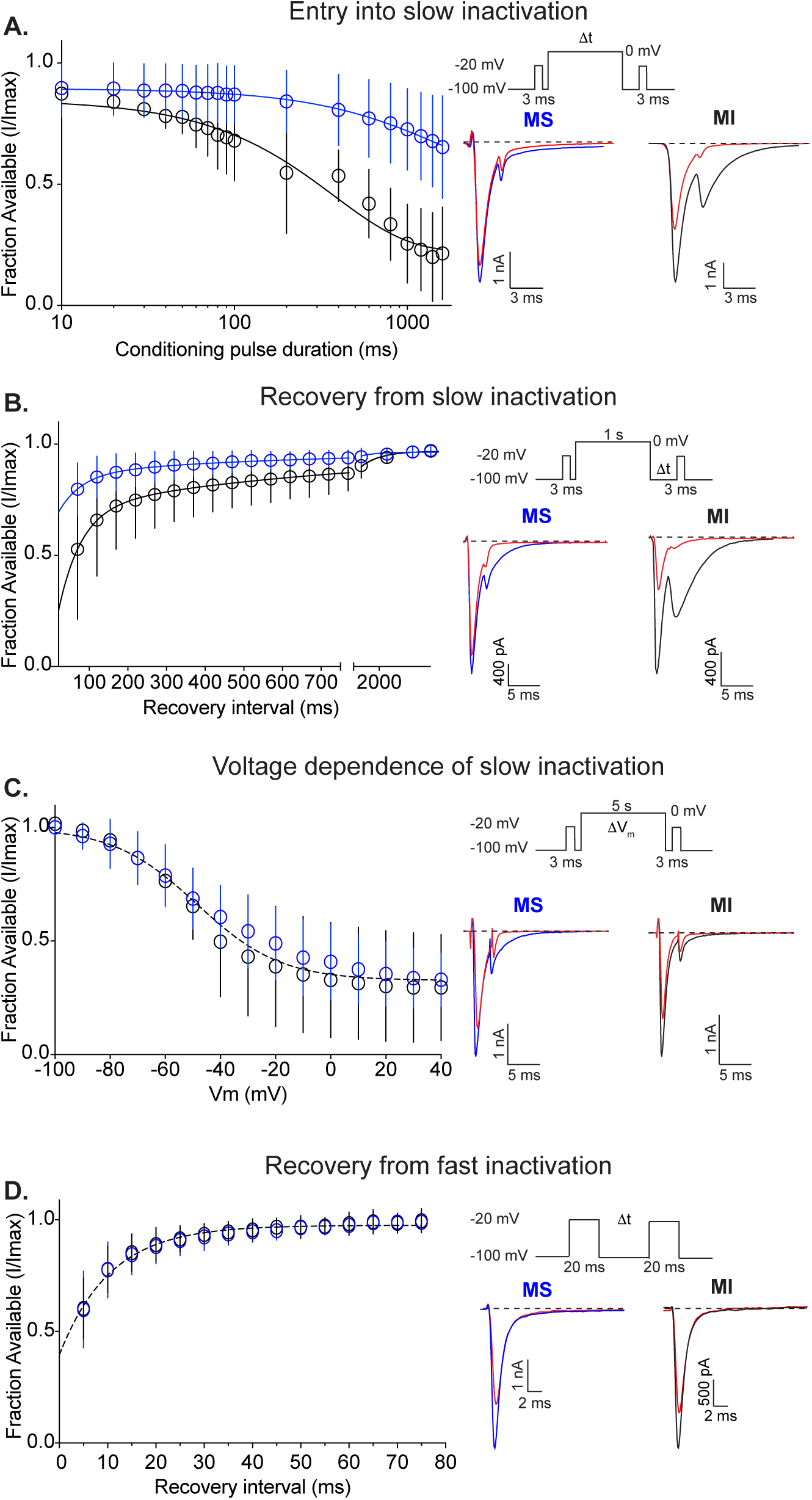
Na_V_ recovery from slow inactivation in small-diameter Vglut3^lineage^ DRG neurons. **A.** Left, Quantification of entry into slow inactivation for menthol-sensitive (MS, blue) and -insensitive (MI, black) Vglut3^lineage^ DRG neurons. The current elicited following a conditioning pulse to 0 mV is normalized to the current elicited during an initial test step and plotted against the duration of the conditioning pulse. Lines show exponential fits to the data. Top right, Voltage protocol used to measure Na_V_ entry into slow inactivation. Channel slow inactivation was elicited by a conditioning pulse ranging from 10-1600 ms. A test step to −20 mV given 12 ms after the conditioning pulse was used to determine entry into slow inactivation. Bottom right, Representative whole-cell voltage-clamp traces of currents elicited from a MS and a MI neurons. Blue or black traces represent initial test steps; red traces represent test pulses given after a 200 ms conditioning step. **B.** Left, Quantification of recovery from slow inactivation kinetics for MS and MI neurons. Recovery during the second test step is normalized to the current during the initial test step and plotted against the recovery interval. Lines show double-exponential fits to the data. MS neurons: n = 9, τ_1_ = 571.1 ms, τ_2_ = 59.5 ms; MI neurons: n = 6, τ_1_ = 764.6 ms, τ_2_ = 58.2 ms; *P* < 0.0001, Extra sum-of-squares F test. Top right, Voltage protocol used to measure Na_V_ recovery from slow inactivation. Slow inactivation was induced by a 1 s conditioning pulse to 0 mV. Recovery was assayed by 3 ms steps to −20 mV with increasing recovery durations, beginning at 50 ms following the conditioning pulse. Bottom right, Representative whole-cell voltage-clamp traces of currents elicited from a MS (blue) and a MI (black) neuron. Blue or black traces represent initial test steps; red traces represent pulses given 50 ms after the conditioning step. **C.** Left, Quantification of steady-state voltage dependence of slow inactivation. Both groups were well fit by a single Boltzmann equation (V_50_ = −48.6 mV, slope factor = −15.6 mV, n = 7–8 neurons per group, *P* = 0.1031, Extra sum-of-squares F test). Top right, Protocol for measuring voltage dependence of slow inactivation. A conditioning pulse of 5 s to membrane potentials between −100 and +40 mV were followed by a 20 ms step to −100 mV and then a test step to −20 mV. Bottom right, Representative traces of currents elicited from a MS (blue) and a MI (black) neuron. Red traces represent test pulses to - 40 mV. **D.** Left, Quantification of recovery from fast inactivation kinetics. Lines show monoexponential fits to the data. Top right, Protocol to measure recovery from fast inactivation. A 20 ms step to −20 mV from −100 mV is followed by varying durations at the recovery potential (−100 mV) before a second test step to −20 mV. MS and MI datasets were well fit by a single monoexponential equation. τ = 10.5 ms, n = 10 for both groups. Bottom right, Representative traces of currents elicited from a MS (blue) and a MI (black) neuron. Red traces represent test pulses 5 ms following the initial test step. Error bars indicate SD; absent error bars are smaller than symbols.

We next analyzed recovery from slow inactivation by delivering a 3-ms test pulse to −20 mV, followed by a 1-s conditioning step from −100 mV to 0 mV, and a second 3-ms test pulse given at recovery intervals of increasing duration. Recovery time constants from both populations were well-fit with double-exponential functions. Consistent with the slow inactivated state being unstable in menthol-sensitive Vglut3^lineage^ neurons, Na_V_ currents in these cells recovered faster from slow inactivation than those in menthol-insensitive neurons (Figure 4B); however, note that the 1-s conditioning pulse did not drive all channels into the slow inactivated state. Sodium currents in menthol-sensitive Vglut3^lineage^ neurons recovered from slow inactivation with an average weighted time constant of 244.3 ms, whereas menthol-insensitive Vglut3^lineage^ neurons recovered slower, with an average weighted time constant of 311.2 ms (*P* < 0.0001, Extra sum-of-squares F test). Indeed, after 50 ms, ∼80% of the Na_V_ current had recovered in menthol-sensitive neurons, compared with ∼50% in menthol insensitive neurons. The steady-state voltage-dependence of slow inactivation was comparable in menthol-sensitive and -insensitive Vglut3^lineage^ neurons (Figure 4C). These data suggest that in menthol-sensitive neurons, slow inactivated states are less stable across membrane voltages. Finally, recovery from fast inactivation was not distinguishable between menthol-sensitive and -insensitive Vglut3^lineage^ neurons (Figure 4D).

Collectively, these data indicate that the slow inactivated state of Na_V_s expressed in menthol-sensitive Vglut3^lineage^ neurons is unstable, which could explain the capacity of these neurons to sustain action potential firing for prolonged periods of time. Moreover, the kinetics of slow inactivation we obtained for Na_V_ currents in this population of small-diameter neurons suggest they do not rely upon TTX-resistant Na_V_s, which readily enter into the slow inactivated state (Blair and Bean, 2003; Choi et al., 2007).

### Na_V_ expression profiles in small-diameter Vglut3^lineage^ DRG neurons

Given the functional differences in membrane excitability and Na_V_ currents between menthol-sensitive and -insensitive Vglut3^lineage^ neurons, we next asked whether these two populations have distinct expression profiles of Na_V_α subunits, nine of which are encoded in the mammalian genome (Na_V_1.1-Na_V_1.9; Catterall, 2012). To do so, we performed single-molecule multiplex *in situ* hybridization experiments (Figure 5). Menthol-sensitive Vglut3^lineage^ neurons were identified based on TRPM8 mRNA expression. Similarly sized, small-diameter Vglut3^lineage^ neurons lacking TRPM8 expression were considered menthol-insensitive neurons.

**Figure 5.**
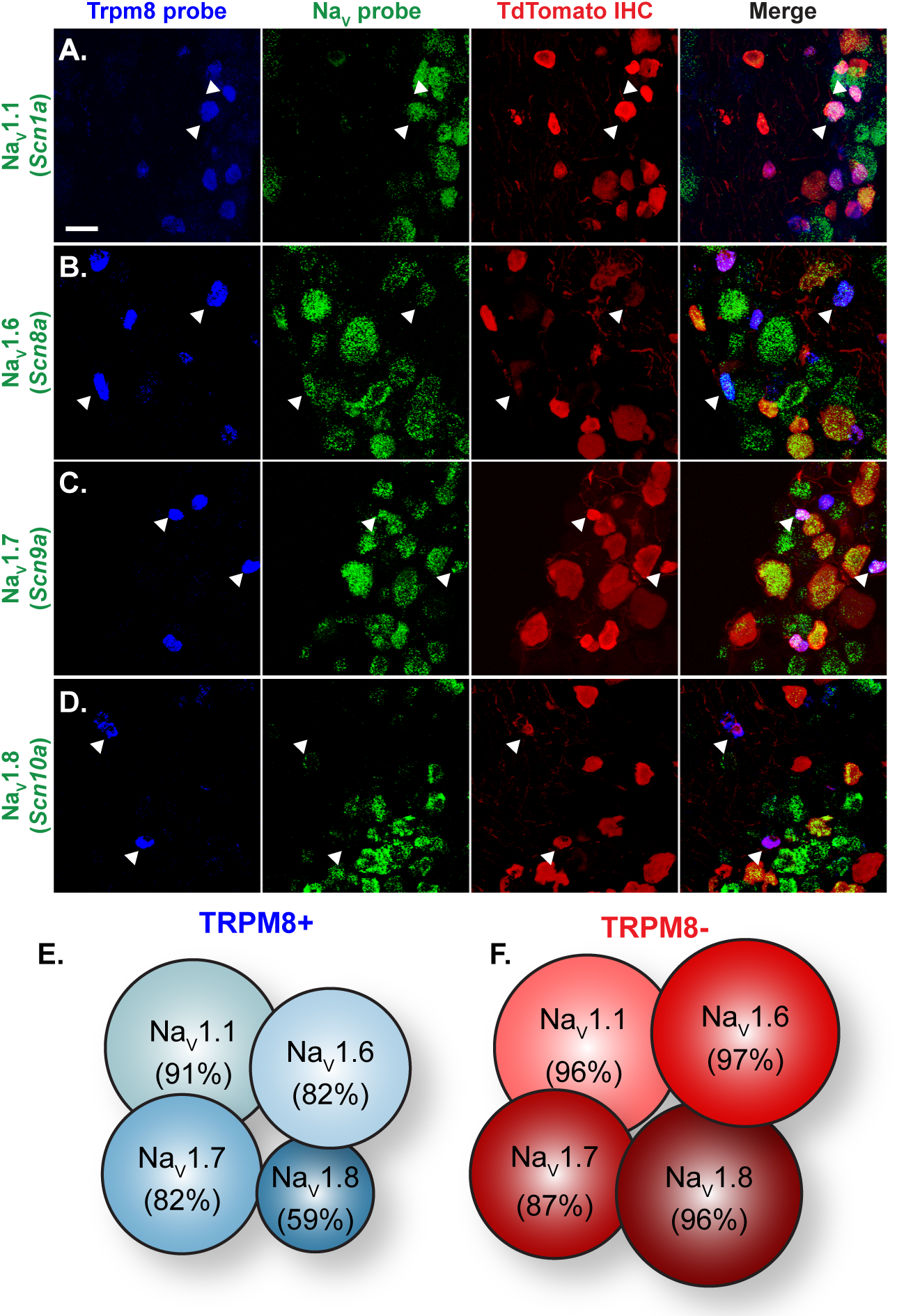
Na_V_ expression profile of small-diameter Vglut3^lineage^ DRG neurons. (**A-D**). Representative confocal images of single molecule multiplex *in situ* hybridizations performed on cryosections of adult DRG (25 μm). Images were acquired with a 40x, 1.3 NA oil immersion objective. Scale bar, 50 μM. Sections were hybridized with probes targeting TRPM8 (*Trpm8*, blue) and the following voltage-gated sodium channel subunits (green): (**A**) Na_V_1.1 (*Scn1a*), (**B**) Na_V_1.6 (*Scn8a*), (**C**) Na_V_1.7 (*Scn9a*), and (**D**) Na_V_1.8 (*Scn10a*). Sections were stained using immunohistochemistry with anti-dsRED (TdTomato, red) to label Vglut3^lineage^ neurons. White arrowheads indicate representative TRPM8+/Na_V_+ neurons. (**E-F**). Schematic representation of the percentage of TRPM8+ (**E**) or TRPM8-(**F**) small-diameter Vglut3^lineage^ neurons that co-labeled for each given Na_V_ subunit.

We focused our analysis on Na_V_1.1, Na_V_1.6, Na_V_1.7 and Na_V_1.8 subunits, which are commonly found in adult murine DRG neurons (Figure 5A-D; Black et al., 1996; Ho and O’Leary, 2011). Quantification of Na_V_ mRNA staining from 848 DRG neurons (n = 3 animals) revealed broad and comparable expression of TTX-sensitive Na_V_1.1, Na_V_1.6, and Na_V_1.7 subunits between TRPM8^+^ and TRPM8^-^ small-diameter Vglut3^lineage^ neurons (Figure 5E-F). Transcripts for the TTX-resistant isoform Na_V_1.8, although widely expressed, was lower in TRPM8^+^ compared with TRPM8^-^ Vglut3^lineage^ neurons, [59% (47/80) vs. 96% (97/101), respectively]. These data suggest that differential expression of Na_V_s at the mRNA level cannot account for the differences in excitability observed between menthol-sensitive and -insensitive Vglut3^lineage^ DRG neurons.

### TTX-sensitive Na_V_s mediate action potential firing in menthol-sensitive Vglut3^lineage^ neurons

Considering the overlap in Na_V_ mRNA expression between putative menthol-sensitive and -insensitive neurons, we next used a pharmacological approach to assess the complement of functional Na_V_ isoforms in these two populations. We first asked whether evoked firing from menthol-sensitive Vglut3^lineage^ neurons is blocked by TTX (Figure 6A). A 1-min application of TTX (0.3 or 1 µM) abolished action potential firing in menthol-sensitive Vglut3^lineage^ neurons (Figure 6B). On the other hand, 1 µM TTX abolished action potential firing in only 3/11 menthol-insensitive Vglut3^lineage^ neurons. The inhibitory effect of TTX on action potential firing was significantly greater in menthol-sensitive neurons compared with menthol-insensitive neurons *(P* = 0.0053, unpaired Student’s *t* test; Figure 6B). Thus, these results demonstrate that menthol-sensitive and -insensitive Vglut3^lineage^ neurons have functionally distinct complements of Na_V_ subunits, with TTX-sensitive channels driving action potential firing in the former and TTX-resistant channels playing a major role in spike firing in the latter.

**Figure 6.**
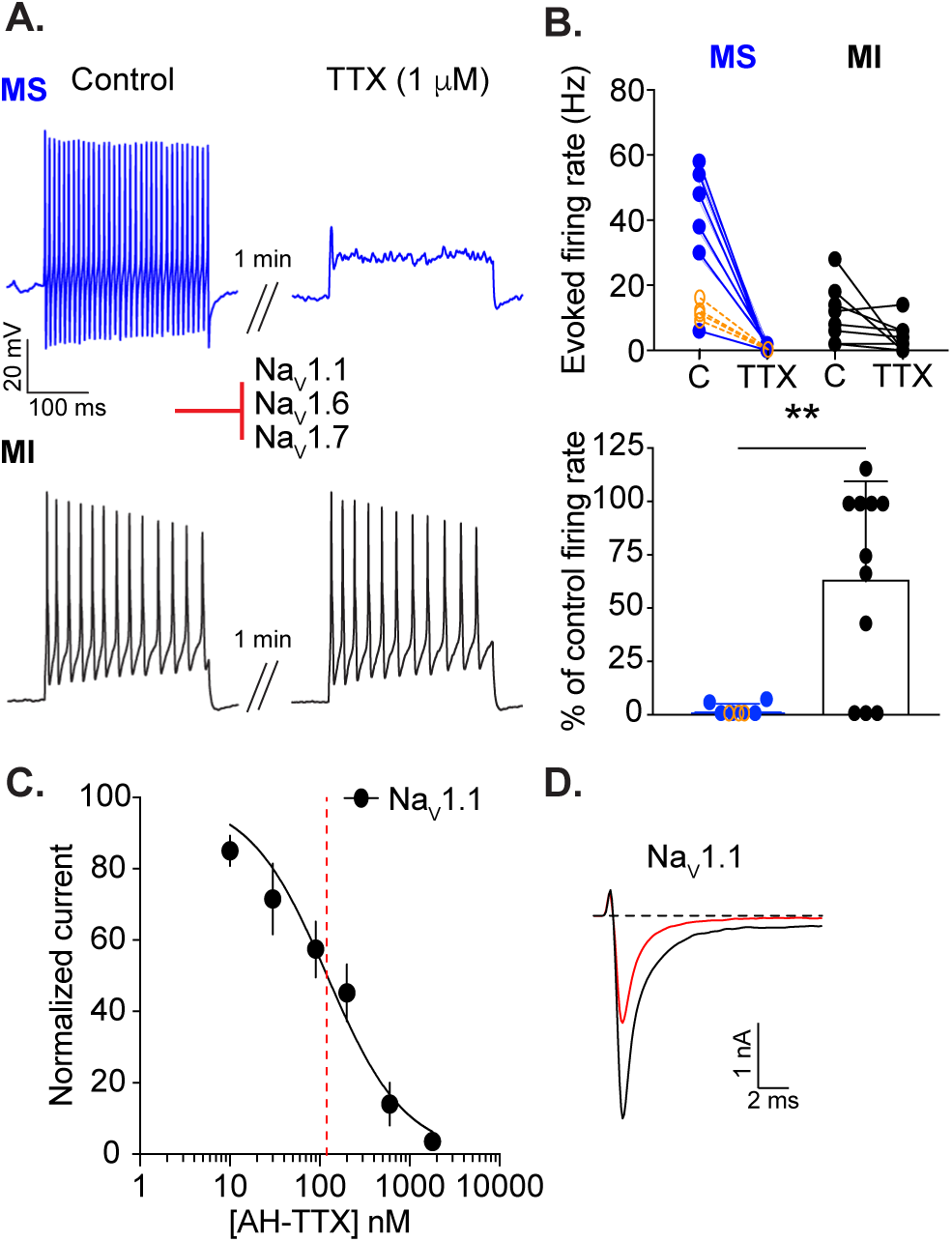
TTX-sensitive Na_V_s mediate action potential firing in menthol-sensitive Vglut3^lineage^ neurons. **(A)** Representative current-clamp traces from menthol-sensitive (top, MS, blue) and -insensitive (bottom, MI, black) Vglut3^lineage^ DRG neurons before (left) and after (right) a 1 min application of TTX (1 μM). **B.** Top, Quantification of firing rates before and after TTX application (n = 6 MS neurons, n = 11 MI neurons). Lines connecting symbols indicate paired observations. Bottom, Quantification of the percentage of control firing rate that remained following TTX application for MS and MI neurons. Blue symbols indicate MS neurons and black symbols indicate MI neurons. Orange symbols indicate firing rates of MS neurons before and after application of 300 nM TTX (n = 4). **C.** A dose-response curve obtained for inhibition of recombinant human Na_V_1.1 channels stably expressed in HEK293 cells by AH-TTX, a blocker of Na_V_1.6 channels. Red dashed line indicates apparent IC_50_. **D.** Representative whole-cell voltage-clamp traces of Na_V_1.1 currents elicited from HEK293 cells. Black trace indicates current elicited prior to application of AH-TTX. Red trace shows reduction in Na_V_1.1 current after application of 200 nM AH-TTX. Red inhibitory sign indicates the inhibition of denoted Na_V_ subunits by the blocker.

We next aimed to dissect the specific contributions of individual TTX-sensitive Na_V_ subunits to action potential firing in menthol-sensitive Vglut3^lineage^ neurons. A metabolite of TTX, 4,9-anhydro-TTX (AH-TTX), has been reported to selectively block Na_V_1.6 channels (Rosker et al., 2007); however, its effect on Na_V_1.1 channels was not examined. Accordingly, we analyzed inhibition by AH-TTX of sodium currents HEK cells stably transfected with human Na_V_1.1 channels (Kahlig et al., 2010). The average peak amplitude for Na_V_1.1 currents recorded from this cell line was −2499 ± 1499 pA (n = 15). The dose-response curve obtained showed an apparent IC_50_ of 120.7 nM (n = 3-5 observations per concentration, Figure 6C). Indeed, there was notable block of Na_V_1.1 currents by 200 nM AH-TTX (Figure 6D), which is within the range of concentrations typically used to block Na_V_1.6 (100–300 nM, Rosker et al., 2007; Hargus et al., 2013; Barker et al., 2017). Thus, we were unable to use this reagent to examine a specific role for Na_V_1.6 channels in action potential firing in menthol-sensitive Vglut3^lineage^ neurons.

### A role for Na_V_1.1 in menthol-sensitive Vglut3^lineage^ neurons

The TTX-sensitive channels Na_V_1.1 and Na_V_1.7 are both expressed in adult DRG neurons and have been implicated in various forms of pain processing (Cummins et al., 2004; Nassar et al., 2004; Osteen et al., 2016). Whether or not they function small-diameter Vglut3^lineage^ DRG neurons has yet to be determined. We therefore investigated the contribution of these subunits to action potential firing in menthol-sensitive neurons.

We first tested ICA 121431, an inhibitor of the Na_V_1.1 and Na_V_1.3 channels (Figure 7A-B). Na_V_1.3 is expressed at only low levels in uninjured adult rat, mouse and human DRGs (Waxman et al., 1994; Felts et al., 1997; He et al., 2010; Usoskin et al., 2015; Chang et al., 2018). Thus, we used ICA 121431 as a blocker of Na_V_1.1 channels in adult mouse DRG preparations (McCormack et al., 2013). Application of 500 nM ICA 121431 drastically reduced action potential firing in menthol-sensitive Vglut3^lineage^ neurons (baseline: 37.1 ± 7.8 Hz, post-ICA 121431: 4.6 ± 4.9 Hz; n = 11, Figure 7B). Action potential firing was also reduced in a subset of menthol-insensitive Vglut3^lineage^ neurons (5.5 ± 4.4 to 2.0 ± 1.9 Hz; n = 8). Like TTX, however, the effect of ICA 121431 was significantly greater in menthol-sensitive compared with menthol-insensitive neurons (*P* = 0.0363, Figure 7B). The specificity of ICA 121431’s block of Na_V_1.1 currents was confirmed by testing inhibition of recombinant Na_V_1.1, Na_V_1.6 and Na_V_1.7 mediated currents (IC_50_ = 25.2 nM, 2.7 µM, and 2.8 µM respectively, n = 4-6 observations per concentration, Figure 7C-D). The average peak currents recorded for Na_V_1.6 and Na_V_1.7 channels were −1545 ± 1134 pA and −1147 ± 1391 pA (n = 6 and n = 9, respectively).

**Figure 7.**
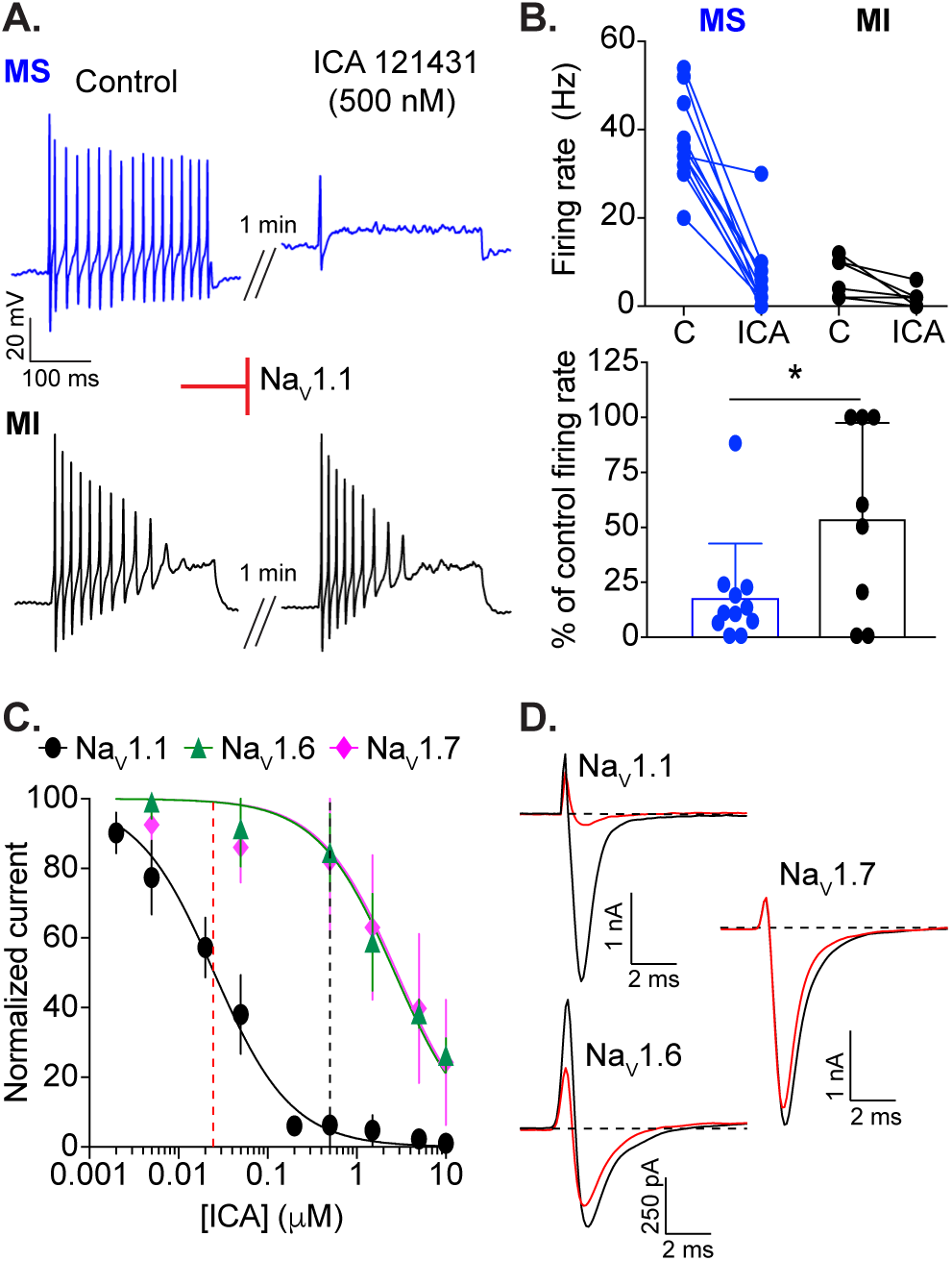
A critical role for Na_V_1.1 channels in action potential firing by menthol-sensitive Vglut3^lineage^ DRG neurons. **(A).** Representative current-clamp traces from menthol-sensitive (top, MS, blue) and -insensitive (bottom, MI, black) Vglut3^lineage^ DRG neurons before (left) and after (right) a 1 min application of ICA 121431 (500 nM). **B.** Top, Quantification of firing rates before and after ICA 121431 application (n = 11 MS neurons, n = 8 MI neurons). Lines connecting symbols indicate paired observations. Bottom, Quantification of the percentage of control firing rate that remained following ICA 121431 application. **C**. A dose-response curve quantifying inhibition of recombinant human Na_V_1.1 (black circles), Na_V_1.6 (green triangles) and Na_V_1.7 (magenta diamonds) by ICA 121431. The apparent IC_50_ for Na_V_1.1 is indicated by a red dashed line. The concentration of ICA 121431 used in this study is indicated by a black dashed line. **D.** Representative whole-cell voltage-clamp traces of Na_V_1.1, Na_V_1.6 and Na_V_1.7 currents elicited from HEK293 cells before (black trace) and after (red trace) application of 500 nM ICA 121431. Red inhibitory sign indicates the inhibition of denoted Na_V_ subunits by the blocker.

We next investigated the contribution of Na_V_1.7 channels to action potential firing in menthol-sensitive neurons using PF 05089771 (25 nM; Alexandrou et al., 2016; Theile et al., 2016), a selective blocker of this channel. PF 05089771 had little effect on mean firing rates in menthol-sensitive Vglut3^lineage^ neurons (Figure 8A-B). Moreover, when cells were analyzed as a percentage of control firing, the effects of PF 05089771 did not differ between menthol-sensitive and menthol-insensitive neurons. A dose-response curve performed in HEK cells transiently transfected with recombinant Na_V_1.7 showed an apparent IC_50_ of PF 05089771 for inhibition of Na_V_1.7 of 10.7 nM (n = 4-5 observations per concentration, Figure 8C), consistent with published values (Alexandrou et al., 2016; Theile et al., 2016). Thus, at the concentration used in this study, PF 05089771 blocked roughly 70% of the Na_V_1.7 mediated current (Figure 8D). We also tested the spider venom toxin Pn3a (300 nM), a structurally unrelated Na_V_1.7 antagonist whose mechanism is distinct from that of PF 05089771 (Deuis et al., 2017). Consistent with results obtained using PF 05089771, Pn3a had no effect on action potential firing in menthol-sensitive neurons (control: 38.7 ± 5.0 Hz, after Pn3a perfusion: 34.0 ± 4.0 Hz, n = 3, Figure 8B). Together, these results demonstrate that action potential firing in menthol-sensitive Vglut3^lineage^ neurons depends upon TTX-sensitive Na_V_s including Na_V_1.1.

**Figure 8.**
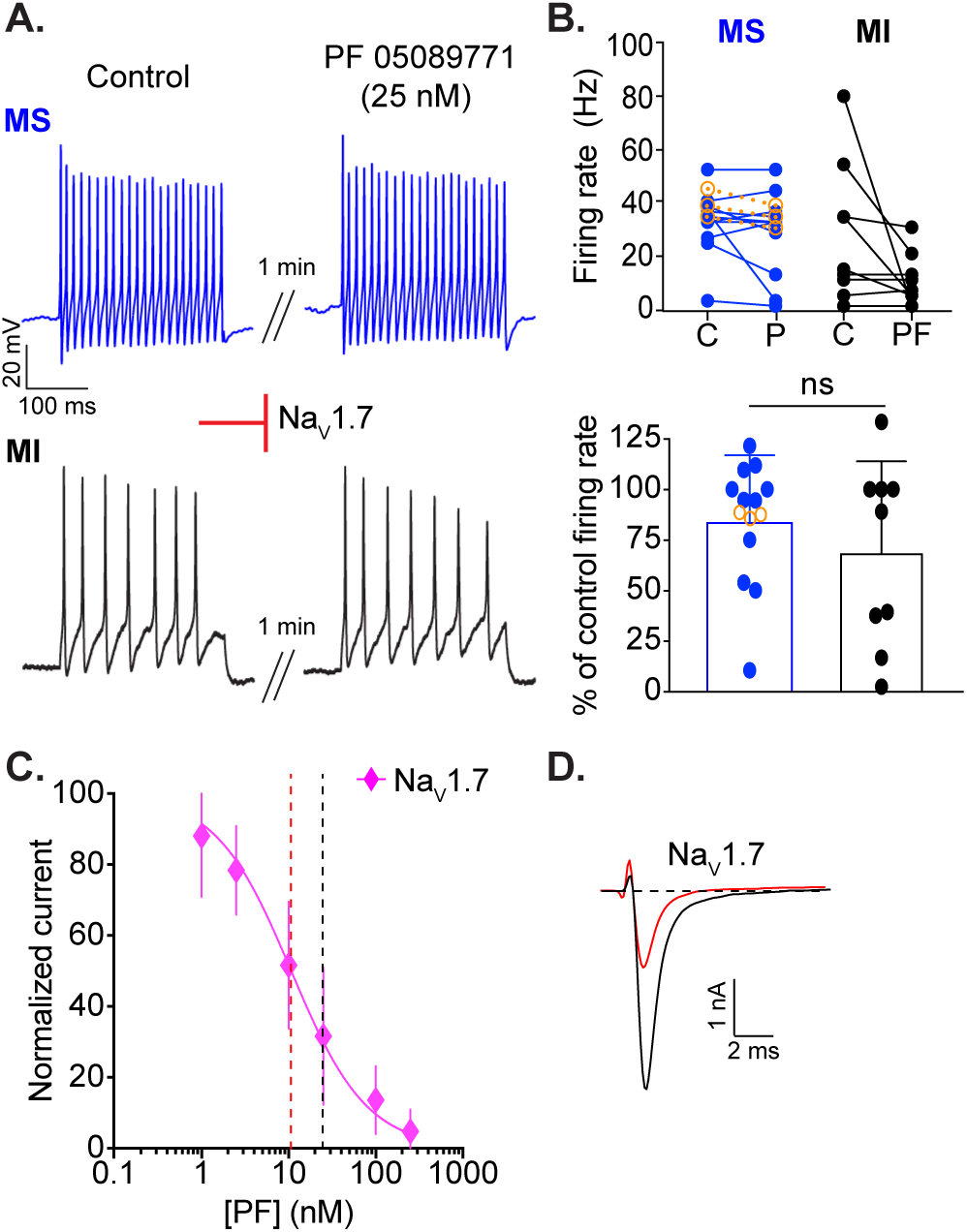
Na_V_1.7 channels do not contribute to action potential firing in small-diameter Vglut3^lineage^ DRG neurons. **(A)**. Representative current-clamp traces from menthol-sensitive (top, MS, blue) and -insensitive (bottom, MI, black) Vglut3^lineage^ DRG neurons before (left) and after (right) a 1 min application of PF 05089771 (25 nM). **B.** Top, Quantification of firing rates before and after PF 05089771 application (n = 11 MS neurons, n = 9 MI neurons) or Pn3a (300 nM, orange symbols in panel F; n = 3 MS neurons). Lines connecting symbols indicate paired observations. Bottom, Quantification of the percentage of control firing rate that remained following PF 05089771 application. **C.** A dose response curve measuring inhibition of recombinant human Na_V_1.7 channels by PF 05089771. The apparent IC_50_ is indicated by a red dashed line. The concentration of PF 05089771 used in this study is indicated by a black dashed line. **D.** Representative whole-cell voltage-clamp traces of Na_V_1.7 currents elicited from HEK293 cells before (black trace) and after (red trace) application of 25 nM PF 05089771. Red inhibitory sign indicates the inhibition of denoted Na_V_ subunits by the blocker. Significance was determined by unpaired Student’s *t* tests. \**P*<0.05, \*\**P*<0.01.

### Na_V_1.1 channels are critical determinants of entry into slow inactivation in menthol-sensitive Vglut3^lineage^ neurons

As Na_V_ currents in menthol-sensitive Vglut3^lineage^ neurons are resistant slow inactivation, we next asked if this biophysical feature depended upon the activity of Na_V_1.1 channels. To accomplish this, we analyzed rates of entry into, and recovery from, slow inactivation in menthol-sensitive Vglut3^lineage^ neurons in the presence of 500 nM ICA 121431. Analysis of whole-cell currents found that the ICA-sensitive component was 38.3% ± 20.2% of the total Na_V_ current in these neurons (Figure 9A-B). In line with Na_V_1.1 channels being critical to the excitability of menthol-sensitive DRG neurons, we found the rate of entry into slow inactivation drastically increased in the presence of ICA 121431. The previously observed rate of 1485 ms fell to 327.7 ms when Na_V_1.1 channels were blocked (Figure 9C; n = 10). Conversely, the average weighted time constant of recovery from slow inactivation more than doubled [without ICA 121431: 311.2 ms, versus with ICA 121431: 686.4 ms (τ_1_ = 725.1 ms, τ_2_ = 59.8 ms), n = 6, Figure 9D]. Linear regression analysis showed a significant correlation between the magnitude of the ICA-sensitive current and the rate of entry into slow inactivation (r^2^ = 0.43, *P* = 0.04, n = 10, Figure 9E). On the other hand, the amplitude of the ICA-sensitive current did not correlate with the rate of recovery from slow inactivation (r^2^ = 0.37, *P* = 0.20, n = 6, Figure 9F). These data demonstrate a new role for Na_V_1.1 in setting the rate of Na_V_ current entry into slow inactivation in sensory neurons. Collectively, our results support a role for Na_V_1.1 channels as key mediators of excitability in menthol-sensitive Vglut3^lineage^ neurons.

**Figure 9.**
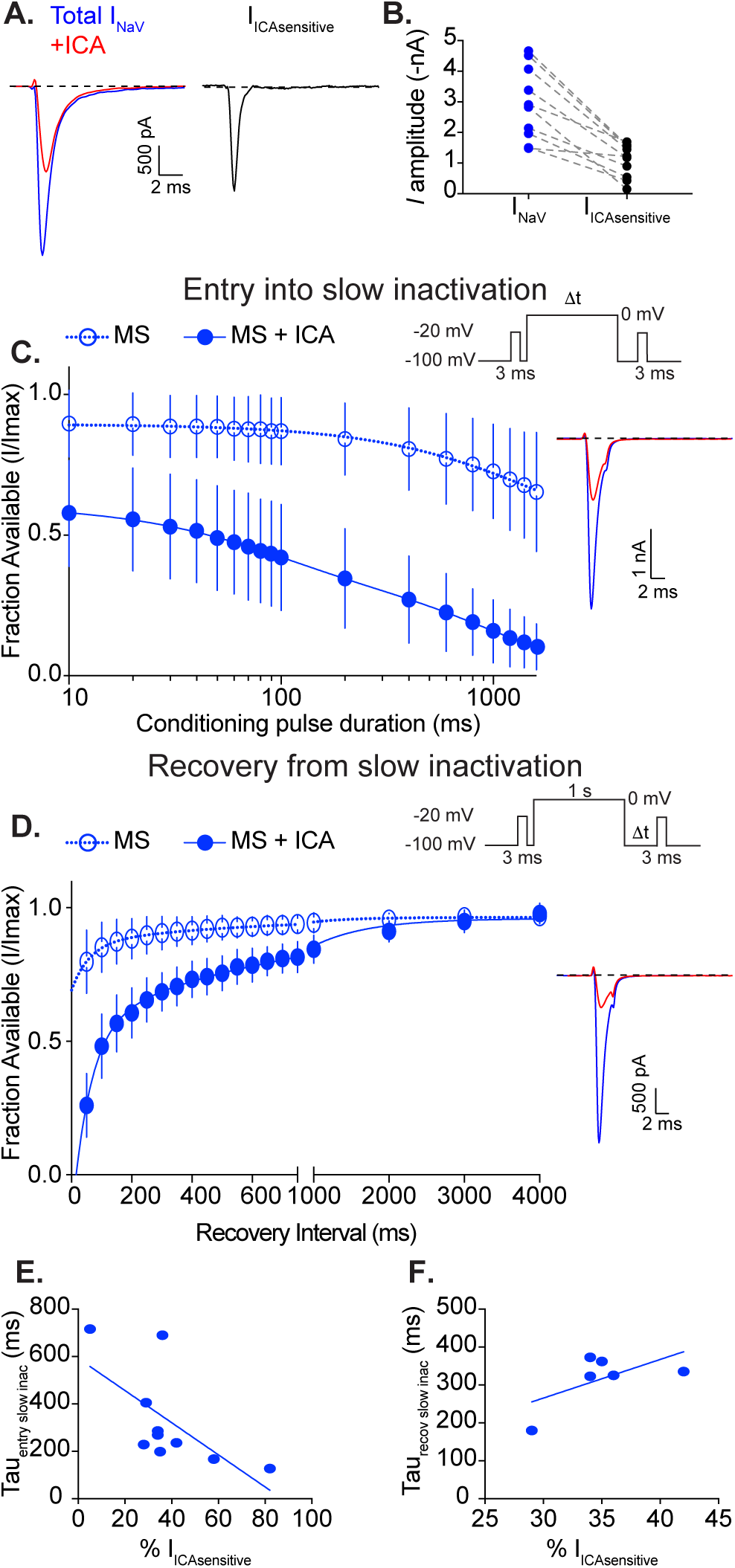
Na_V_1.1 channels determine entry into slow inactivation rates in menthol-sensitive Vglut3^lineage^ neurons. **A.** Representative whole-cell voltage-clamp traces of currents elicited from a MS neuron in the before (blue trace) and after application of 500 nM ICA 121431 (red trace). The subtracted Na_V_1.1-mediated current is shown in black. **B.** Quantification of peak Na_V_ current amplitude before (blue circles) and after ICA 121431 application (black circles). Gray dashed lines indicated paired observations. **C.** Quantification of entry into slow inactivation for MS Vglut3^lineage^ DRG neurons during blockade of Na_V_1.1 channels by ICA 121431 (filled circles, blue lines). Data for MS neurons from Figure 4A is shown for comparison (clear circles, dashed lines). The current elicited following a conditioning pulse to 0 mV is normalized to the current elicited during an initial test step and plotted against the duration of the conditioning pulse. Lines show exponential fits to the data. Top right, Voltage protocol used to measure Na_V_ entry into slow inactivation. Bottom right, Representative whole-cell voltage-clamp traces. Blue trace represents initial test step; red trace represents test pulses given after a 200 ms conditioning step. **D.** Left, Same as C except Quantification represents recovery from slow inactivation kinetics. Data for MS neurons from Figure 4B is shown for comparison (clear circles, dashed lines). Recovery during the second test step is normalized to the current during the initial test step and plotted against the recovery interval. Lines show double-exponential fits to the data. MS neurons + ICA: τ_1_ = 725.1 ms, τ_2_ = 59.8 ms, n = 6. Top right, Voltage protocol used to measure Na_V_ recovery from slow inactivation. Bottom right, Representative whole-cell voltage-clamp trace of current elicited from a MS neuron in the presence of ICA 121431. Blue trace represents initial test step; red trace represents pulse given 50 ms after the conditioning step. **E.** Individual entry into slow inactivation rates for MS neurons plotted against the percentage of the total Na_V_ current that was sensitive to ICA 121431; *r^2^ = 0.43*, *P = 0.04*. **F.** Same as **E** but individual recovery from slow inactivation rates are plotted; *r^2^ = 0.3*7, *P = 0.20*.

## Discussion

Small-diameter Vglut3^lineage^ DRG neurons are a heterogeneous population that encode distinct somatic senses. This study reveals two important findings about the functional heterogeneity in such neurons. First, menthol-sensitive Vglut3^lineage^ DRG neurons possess a unique excitability profile, which allows them to maintain prolonged spike discharges. Second, TTX-sensitive Na_V_s mediate action potential firing in these sensory neurons, with a notable contribution of Na_V_1.1. We propose that cation influx through TRPM8 ion channels produces an excitatory drive that activates Na_V_1.1 ion channels at room temperature. Once activated, these channels cycle through open and fast-inactivated states, with the majority of channels bypassing long-lived slow inactivated states. This is likely attributable to unique features of Na_V_1.1-containing macromolecular complexes in menthol-sensitive neurons, including association with auxiliary proteins or posttranslational modifications (Aman and Raman, 2007), with the end result being continuous action potential firing (Figure 10). Thus, menthol-sensitive Vglut3^lineage^ DRG neurons represent a highly excitable population of small-diameter sensory neurons in which action potential firing depends upon TTX-sensitive Na_V_ complexes.

**Figure 10.**
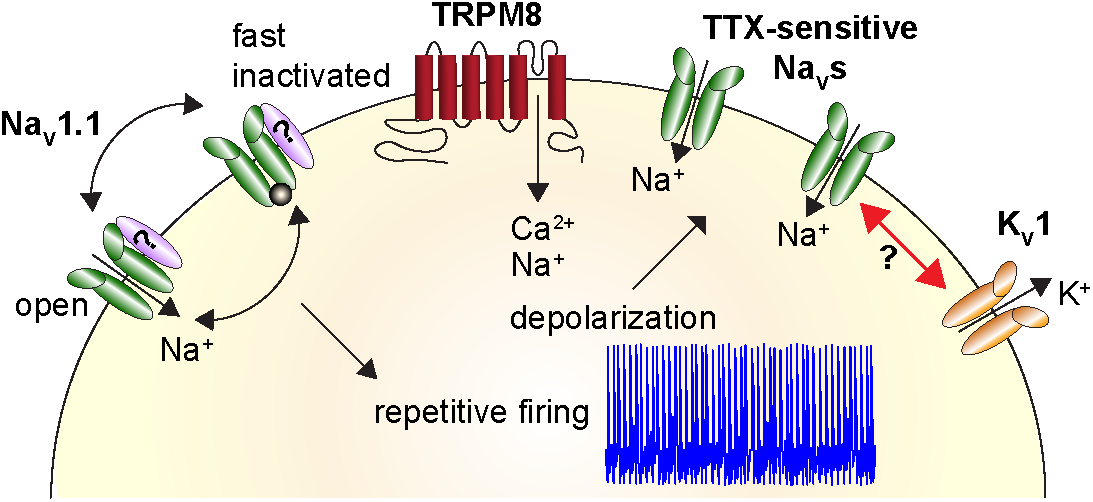
Proposed model demonstrating heightened excitability of menthol-sensitive Vglut3^lineage^ DRG neurons. Activation of TRPM8 ion channels (red) causes an influx of cations that depolarizes the neuron. This leads to activation of TTX-sensitive Na_V_s, including Na_V_1.1 channels (green), and subsequent action potential firing. Repetitive firing is achieved by TTX-sensitive Na_V_s cycling through open and fast inactivated states, without being sequestered into long-lived slow inactivated states. This may be facilitated by association with auxiliary proteins (purple). Na_V_ complexes in menthol-sensitive Vglut3^lineage^ DRG neurons could function to oppose a strong hyperpolarizing conductance mediated by K_V_1 potassium channels (orange).

Prior work has focused on the role of potassium channels as excitability breaks in in TRPM8-expressing sensory neurons. A molecular profiling study identified the TASK-3 leak potassium channel as highly enriched in TRPM8^+^ DRG neurons and suggested that inhibition of this channel decreases cold activation thresholds (Morenilla-Palao et al., 2014). There was only a modest effect, however, of TASK-3 genetic deletion on intrinsic excitability. H-current (I_h_), a conductance mediated by hyperpolarization-activated cyclic nucleotide-gated (HCN) channels, is reportedly pronounced in cold-activated sensory neurons (Viana et al., 2002; Orio et al., 2009). Furthermore, genetic deletion of *Hcn1* converts firing patterns in cold-sensing optic nerve fibers from regular to burst spiking (Orio et al., 2012). Consistent with previous studies, we observed a prominent sag ratio, a current-clamp readout of I_h_, in menthol-sensitive Vglut3^lineage^ neurons; however, our study revealed no difference in sag ratio between menthol-sensitive and -insensitive Vglut3^lineage^ neurons. Thus, TASK-3 and HCN channels, though important for cold detection, are unlikely to mediate the differences in intrinsic excitability between these two populations. Future studies are needed to determine whether other potassium conductances, such as those mediated Kv1 (Madrid et al., 2009; Gonzalez et al., 2017a; Gonzalez et al., 2017b), contribute to differences in intrinsic excitability between menthol-sensitive and - insensitive DRG neurons.

In addition to potassium channels, previous studies investigated TTX-resistant Na_V_s in menthol-sensitive DRG neurons. Na_V_1.8 is found in ∼90% of small-diameter DRG neurons (Shields et al., 2012) and has been implicated in menthol-sensitized cold responses (Zimmermann et al., 2007). However, Na_V_1.8 null mice show normal physiological and behavioral responses to cold (Luiz et al., 2019). A recent study identified a subpopulation of DRG neurons that are both Vglut3^lineage^ and Na_V_1.8^lineage^ (Patil et al., 2018). These neurons possess properties to similar the menthol-sensitive neurons analyzed in the present work, including fast action potential durations, insensitivity to capsaicin, and small somata. Moreover, Na_V_1.9, the other TTX-resistant Na_V_ isoform, was reported to be expressed in nociceptors that respond to cooling, as well as contribute to pain perception in response to noxious cold (Lolignier et al., 2015). In that study, however, Na_V_1.9 mRNA co-expressed with only ∼20% of TRPM8^+^ DRG neurons. The proportion of adult menthol-sensitive neurons that express functional Na_V_1.8 or Na_V_1.9 protein is unknown; nonetheless, our pharmacological studies indicate that TTX-resistant Na_V_s do not drive action potential firing in menthol-sensitive neurons under our experimental conditions.

Instead, we provide evidence that functionally distinct Na_V_s contribute to the different excitability profiles of menthol-sensitive and -insensitive Vglut3^lineage^ DRG neurons. Although multiplex *in situ* hybridization data showed widespread expression of several Na_V_ transcripts, our pharmacological analysis revealed a more restricted functional contribution, with Na_V_1.1 comprising over one third of the total Na_V_ current and mediating most action potential firing in menthol-sensitive Vglut3^lineage^ DRG neurons. Conversely, TTX-resistant channels dominated in menthol-insensitive neurons. In DRG, Na_V_1.1 is reported to be predominantly localized to medium-diameter neurons that mediate mechanical pain (Osteen et al., 2016). Interestingly, that study showed that ∼40% of trigeminal neurons that express functional Na_V_1.1 channels also exhibit menthol-evoked calcium transients. Our study extends these findings by demonstrating that action potential firing in menthol-sensitive Vglut3^lineage^ DRG neurons is dependent upon TTX-sensitive Na_V_s, with the Na_V_1.1/Na_V_1.3 antagonist ICA 121431 dramatically reducing firing rates. Furthermore, while TTX-sensitive Na_V_1.7 channels are important to the function of small-diameter nociceptors and pain signaling (Cox et al., 2006; Minett et al., 2012; Yang et al., 2018), these channels are likely inactivated at the resting membrane potential of menthol-sensitive neurons. Indeed, the V_1/2_ of inactivation of Na_V_1.7 is roughly −75 mV (Alexandrou et al., 2016). Conversely, the V_1/2_ inactivation of Na_V_1.1 is approximately −17 mV (Aman et al., 2009), a membrane potential that is much more depolarized than the resting potential of menthol-sensitive Vglut3^lineage^ DRG neurons in our study. Thus, we have identified a new role for TTX-sensitive Na_V_1.1 channels in action potential firing in small-diameter DRG neurons.

Na_V_1.1 channels promote excitability and high frequency firing in several neuronal populations. In mouse models of irritable bowel syndrome and chronic visceral hypersensitivity, Na_V_1.1 is functionally upregulated, leading to hyperexcitability of mechanosensory fibers innervating the colon (Osteen et al., 2016; Salvatierra et al., 2018). Moreover, mutations in Na_V_1.1 are most frequently associated with inherited forms of epilepsy, including Dravet syndrome (Catterall et al., 2010). In this disorder, loss of Na_V_1.1 in hippocampal interneurons leads to reduced sodium current and attenuated action potential firing (Yu et al., 2006), resulting in disinhibition of hippocampal circuits that causes seizures (Oakley et al., 2013). To our knowledge, our results provide the first functional evidence for Na_V_1.1-dependent action potential firing in small-diameter somatosensory neurons.

Our data also indicate that the biophysical properties of Na_V_1.1-containing channel complexes could explain the heightened excitability of menthol-sensitive DRG neurons. Na_V_ currents in these neurons entered into slow inactivation much more slowly than what has been reported for other DRG populations (Blair and Bean, 2003), with a time constant of ∼1.5 s (Figure 4A). This contrasts with capsaicin-sensitive nociceptors and IB_4_^+^ DRG neurons, where slow inactivation of TTX-resistant Na_V_s is reported to produce action potential adaptation in response to sustained depolarization (Blair and Bean, 2003; Choi et al., 2007). Importantly, application of ICA 121431 drastically enhanced the rate of entry into slow inactivation in menthol-sensitive DRG neurons (Figure 9C). Thus, the resistance of Na_V_1.1 currents to slow inactivation could be a mechanism by which menthol-sensitive neurons sustain action potential firing for extended periods of time.

Previous studies have reported that Na_V_1.1 channels are subject to use-dependent inactivation at high firing frequencies (Spampanato et al., 2001). It is therefore possible that in menthol-sensitive neurons, Na_V_1.1 α subunits associate with auxiliary proteins that destabilize inactivated states, such as the β4 subunit (Aman et al., 2009). Moreover, due to the lack of selective pharmacological tools, we were unable to test the contribution of Na_V_1.6 channels to action potential firing in small-diameter Vglut3^lineage^ DRG neurons. It has been hypothesized, however, that synergistic activity of Na_V_1.1 and Na_V_1.6 is important for overcoming the high action potential threshold set by voltage-gated potassium channels of the K_V_1 family in pyramidal cells and GABAergic interneurons (Lorincz and Nusser, 2008). K_V_1 channels are also expressed in TRPM8^+^ trigeminal neurons, where they are proposed to determine thermal excitability (Madrid et al., 2009). We cannot rule out the possibility that action potential firing patterns in menthol-sensitive Vglut3^lineage^ DRG neurons are tuned by the concerted actions of Na_V_1.1 and Na_V_1.6 channels that counterbalance an opposing Kv1 conductance, thus regulating the responsiveness of these neurons (Figure 10).

The finding that menthol-sensitive neurons are a subset of Vglut3^lineage^ neurons raises the possibility that Vglut3 protein plays a role in synaptic transmission from TRPM8-expressing DRG neurons to second order neurons in the spinal cord. In contrast to this model, Vglut3^-/-^ mice are reported to have normal responses to cold stimuli, indicating that Vglut3 protein is not required for TRPM8-dependent behaviors in mice (Draxler et al., 2014). Furthermore, in sensory neurons innervating the dura and cerebral blood vessels, TRPM8 and Vglut3 protein expression do not overlap in adult mice (Ren et al., 2018). Thus, we speculate that menthol-sensitive neurons express the *Slc17A8* locus during development rather than in mature DRG.

Collectively, our data indicate that, unlike many other small-diameter DRG populations, action potential firing in in menthol-sensitive DRG neurons is dependent upon TTX-sensitive Na_V_s including Na_V_1.1. Genetic approaches using Na_V_1.1 null mutations are needed to define the exact contributions of this subunit to the function of menthol-sensitive neurons, as well as sensory-driven behaviors (Cheah et al., 2012). It also remains to be determined if Na_V_1.1 channels are viable therapeutic targets for pathologies that produce cold hypersensitivity. Additionally, menthol has been used for centuries as a topical analgesic and anti-pruritic. Indeed, it has been shown that TRPM8-expressing DRG neurons are required for inhibition of itch by cooling and furthermore, that topical application of menthol inhibits chloroquine-evoked itch behaviors (Palkar et al., 2018). Thus, targeting TTX-sensitive Na_V_1.1 channels in menthol-sensitive DRG neurons might prove to be a new direction for the treatment of various sensory disorders.

## Acknowledgements

This research was supported by NIAMS R01AR051219 (to E.A.L.). T.N.G. holds a Postdoctoral Enrichment Program Award from the Burroughs Wellcome Fund and was supported by NHLBI T32HL120826. Core facilities were supported by the Columbia University EpiCURE Center (NIAMS P30AR069632) and the Thompson Family Foundation Initiative in CIPN and Sensory Neuroscience. This project was initiated during the MBL Neurobiology Course with support from NINDS R25NS063307. Dr. Blair Jenkins, Mr. Javier Marquina-Solis and Dr. Adrian Thompson participated in preliminary studies at MBL. Thanks to Dr. Manu Ben-Johny and Dr. Lori Isom for sharing reagents, Dr. Irina Vetter for peptide toxins, Ms. Venesa Cuadrado for technical assistance, Ms. Rachel Clary for assistance with custom MATLAB routines, and Dr. Jon Sack and members of the Lumpkin laboratory for helpful discussions.

## Author contributions

TNG and EAL performed calcium imaging and data analysis. TNG designed and preformed all electrophysiological experiments and data analysis. TAD assisted TNG with *in situ* hybridization and TAD performed quantitative analysis. TNG made the figures and wrote the first draft of the manuscript. EAL assisted with writing the manuscript. TNG and EAL edited the manuscript and all authors approved the manuscript. EAL and TNG acquired funding and EAL supervised the project.

